# High-throughput experimental validation of novel hairpin ribozymes

**DOI:** 10.1101/2025.07.08.663751

**Authors:** Kaitlyn Matteo, Eric J. Hayden

**Affiliations:** Biomolecular Sciences Graduate Programs, Boise State University, Boise, Idaho 83725, USA; Department of Biological Sciences, Boise State University, Boise, Idaho 83725, USA

**Keywords:** Deep-sequencing, High-throughput, Ribozyme, RNA, Self-cleavage

## Abstract

The small self-cleaving hairpin ribozyme has served as a model for RNA structure and function and has been engineered for biotechnology applications. Hairpin ribozymes were thought to be rare with only four known examples, which limited the interpretation of their biological importance and the starting sequences for engineering efforts. Recently, a bioinformatics approach identified hundreds of different RNA sequences in metatranscriptomic data that matched a novel permutation of the hairpin ribozyme. However, the self-cleavage activity of most of these sequences has not been experimentally demonstrated. Here, a high-throughput sequencing-based approach was used to evaluate the co-transcriptional self-cleavage activity of 855 different hairpin ribozymes in parallel. The results showed that nearly all sequences are very efficient self-cleaving ribozymes, and even rare nucleotides at highly conserved positions did not prevent observable ribozyme activity. The distribution of activity observed suggests that the metatranscriptomic sequences could contain random mutations from efficient wild-type ribozymes. The results further validate the bioinformatics approach that was used for ribozyme discovery and opens further questions about the biological roles of these ribozymes in the diverse environments where they were discovered.

## Introduction

There are several known classes of small self-cleaving ribozymes that catalyze the site-specific hydrolysis of a phosphodiester linkage (Jimenez et al., 2015). Self-cleaving ribozymes are found widely distributed in viruses and organisms spanning the tree of life. They are known to carry out biological roles in rolling-circle replication, retro-transposition, translation initiation and allosteric response systems (Doudna & Cech, 2002; Jimenez et al., 2015). New self-cleaving ribozymes are still being discovered, and their distributions and biological roles are still being investigated (Jimenez et al., 2015; Liu et al., 2021; Ruminski et al., 2011; Weinberg et al., 2021). The hairpin ribozyme is one structural class of self-cleaving RNA molecules that has been studied extensively as a model system for RNA folding and catalysis (Berzal-Herranz et al., 1993; Fedor, 2000; Kath-Schorr et al., 2012; Wilson et al., 2001). Based on an early discovery and structural understanding (Buzayan et al., 1986; Rupert et al., 2002; Rupert & Ferré-D’Amaré, 2001), hairpin ribozymes have also been engineered for biotechnology applications. This includes modification with aptamer sequences to create allosteric “aptazymes”, and other alterations to facilitate spliceosome-like activity and recombinase activity (Hieronymus & Müller, 2021; Vauléon & Müller, 2003; Zhu et al., 2023). Until recently, only four naturally occurring examples of hairpin ribozymes were known (Buzayan et al., 1986; Kalvari et al., 2018; Kaper et al., 1988; Rubino et al., 1990). This rarity suggested that hairpin ribozymes had limited biological impact, and the lack of structural variants limited engineering efforts.

A recent bioinformatics effort identified ∼1000 different hairpin ribozyme sequences (Weinberg et al., 2021). These newly discovered ribozymes were found in metatranscriptomic datasets where many appear to be involved in the replication of virus-like, circular single-stranded RNA molecules. These hairpin ribozymes were discovered using pattern-based computational search methods (DARN! and Infernal) (Weinberg et al., 2021). This approach enabled searching for a novel permutation of hairpin ribozyme, which turned out to be more abundant in these datasets. Several representative sequences from this search were tested previously and were shown to self-cleave with high efficiency. However, most of the discovered sequences remained experimentally uncharacterized. Here, we sought to experimentally evaluate the catalytic activity of the majority of these novel hairpin ribozyme sequences (**Figure 1A**). We chose a diverse sample of hairpin ribozymes (“extra-spruce-initial alignment”) that contained predicted ribozymes with both sequence and length variability. To study this many sequences, we used a high-throughput approach where DNA templates for all the different sequences were pooled together and transcribed simultaneously. The catalytic activity was determined by deep-sequencing to count the number of instances each sequence was observed as self-cleaved or uncleaved during the *in vitro* transcription reaction (**Figure 1B**) (Bendixsen et al., 2019; Kobori & Yokobayashi, 2016; Roberts et al., 2023).

**Figure 1.**
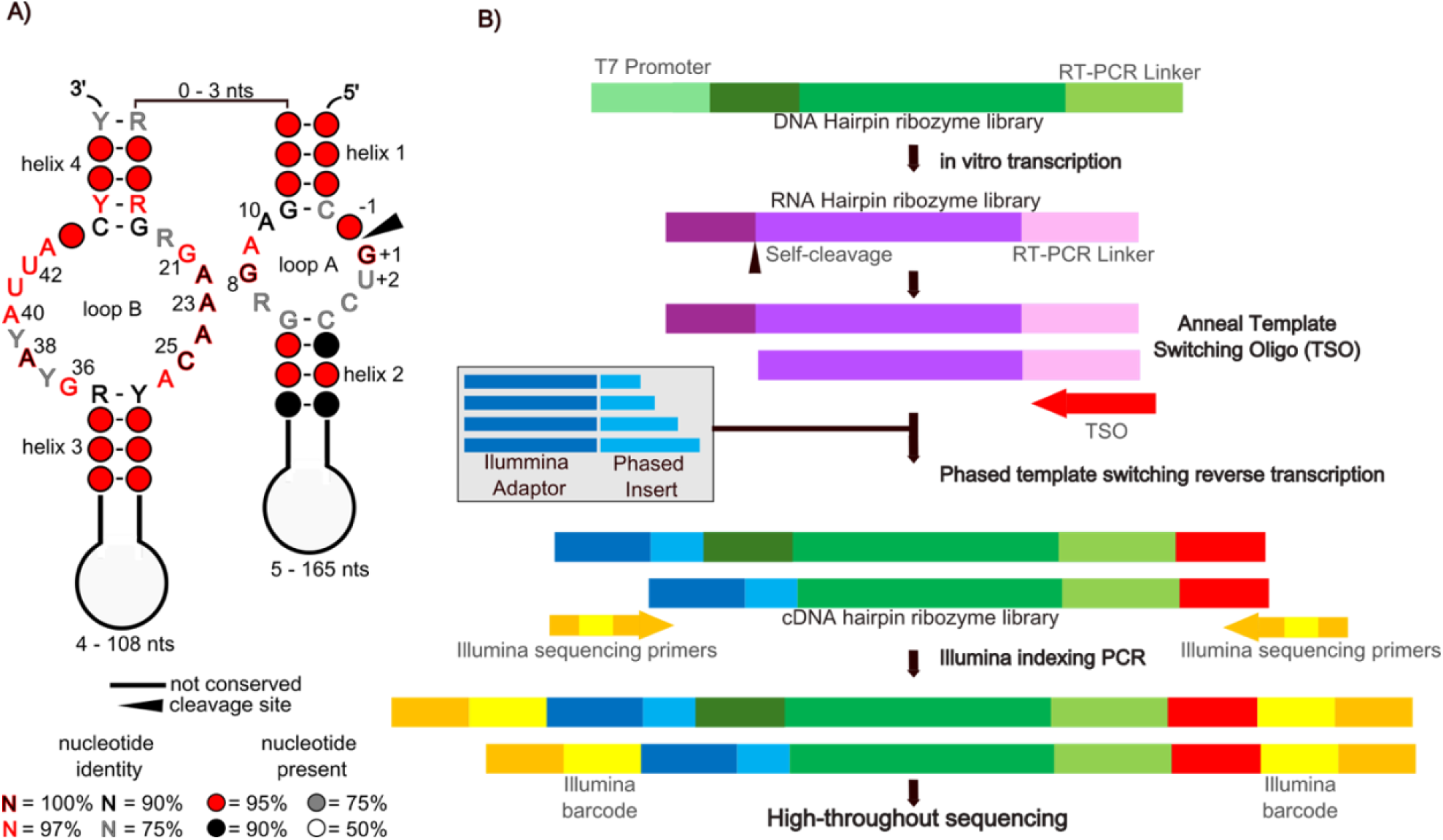
**(A)** Consensus diagram of hairpin ribozymes used in this study. The secondary structure is as previously reported for this permutation of the ribozyme. Nucleotide identities, conservation (see legend), and lengths of variable regions are representative of the 898 ribozymes selected for high-throughput experimental characterization. **(B)** Illustration of the high-throughput co-transcriptional self-cleavage assay.

## Results and Discussion

The high-throughput approach to characterize the cleavage activity of the RNA sequences started with the synthesis of a single-stranded DNA (ssDNA) oligonucleotide pool (IDT) that contained 898 unique sequences, all preceded by a T7 promoter to enable *in vitro* transcription. All sequences ended with the same 43 nucleotide (nt) primer binding sequence for RT-PCR (**Figure 1B**)^22^. Due to length limitations of ssDNA oligo synthesis not all sequences from the reported dataset (“extra-spruce-initial alignment”) were synthesized, and ribozymes exceeding 234 nucleotides (nts) were excluded. The pool was made double stranded by PCR amplification. Transcription reactions were carried out at 37ºC. The reactions contained 2 mM rNTP’s, 30 mM MgCl_2_ and were terminated after one hour with the addition of EDTA to suppress Mg^2+^-dependent activity and placed on ice.

*In vitro* transcribed RNA was reverse-transcribed with a template switching 5’-RACE protocol to convert both cleaved and uncleaved RNA to complementary DNA (cDNA) and add a common primer binding sequence to the variable 5’-end of the RNA (Bendixsen et al., 2019; Kobori et al., 2015). This enabled both cleaved and uncleaved sequences to be amplified by PCR with the same primer pair. The template switching oligos used during 5’-RACE also contained a “phased adaptor” sequence that introduces four different-length sequences before the first nucleotide of the ribozyme, creating balanced nucleotide diversity that improves amplicon sequencing on Illumina platforms (Bendixsen et al., 2020). Illumina dual-index PCR primers (IDT) were used to enable multiplexing of replicates prior to deep-sequencing on an Illumina Miseq.

The relative activity of each RNA sequence was determined by bioinformatics analysis of the MiSeq data using custom python scripts (GitLab). Briefly, sequencing reads were matched to the original pool of references along with its cleavage status. All ribozymes have the same predicted cleavage site near the 5’-end, which enabled identification of each unique RNA sequence, even after the short 5’-end (5 nt) was cleaved off (**Figure 1A**). Cleavage status was determined based on the presence or absence of the short 5’-end sequence upstream of the expected cleavage site. To be counted as uncleaved, sequences were required to match at least three of the five nucleotides of the 5’-fragment and a 100% match to the corresponding 3’ ribozyme sequence. Only sequences ending at the expected self-cleavage site were considered “cleaved”. All other sequences were discarded from further analysis.

To evaluate the relative catalytic activity of the sequences, the number of reads that matched a sequence and ended at the expected self-cleavage site were counted (*N*_*cl*_). The number of reads that matched the uncleaved state (*N*_*uncl*_) was also counted. The fraction cleaved (FC) was calculated as follows:

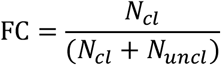

Because all sequences were transcribed at the same time, under the same biochemical conditions, we interpret the FC as an indicator of relative catalytic efficiency that encompasses both folding and catalysis (Kobori et al., 2015; Long & Uhlenbeck, 1994). Co-transcriptional self-cleavage was carried out in triplicate and the mean probability FC value was determined for each sequence. The standard error (SE) and 95% confidence intervals were determined using binomial statistics based on the total number of observations for each individual sequence (**Supplemental_Table_S1**).

### Matteo and Hayden

Our deep-sequencing yielded 1,172,231 reads that passed quality control and were used to determine the FC values for the sequences (**Supplemental_Table_S1)**. To evaluate the range of self-cleaving activities within the studied sequences, we binned sequences by FC and counted the number of sequences that fell into each bin (**Figure 2)**.

**Figure 2.**
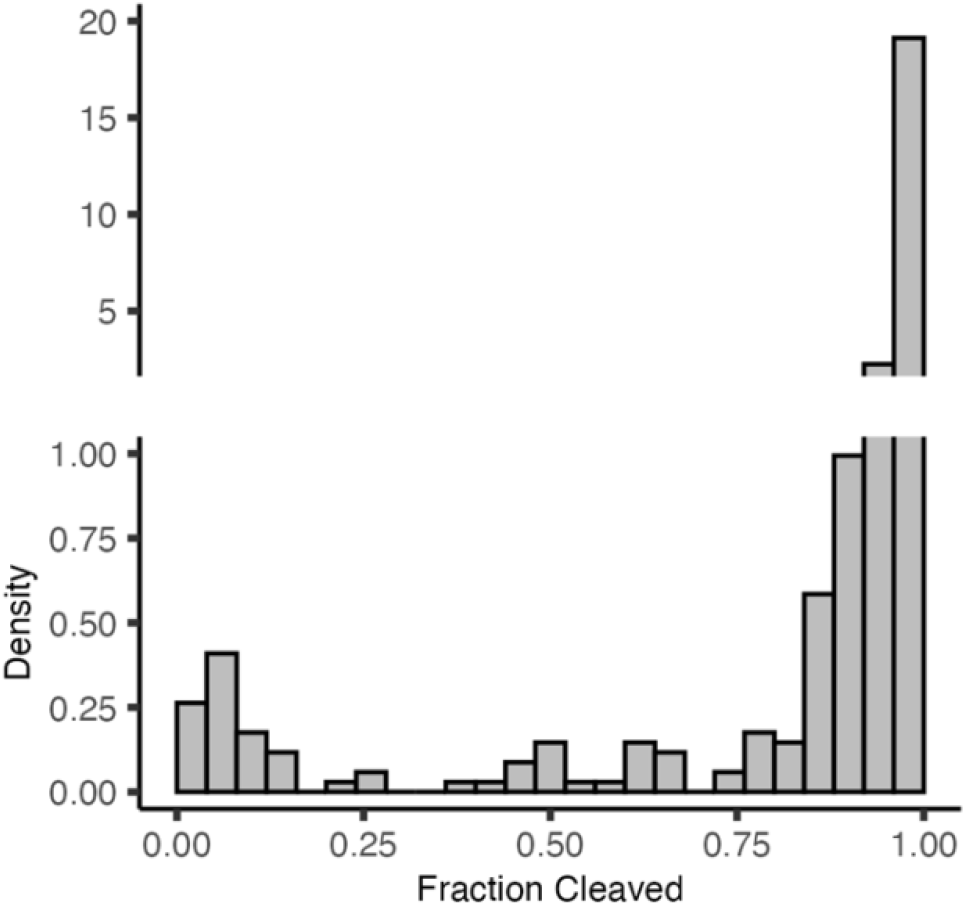
Histogram density plot of fraction cleaved values for the experimentally characterized ribozymes. Ribozymes were binned in increments of 0.04 for a total of 25 bins, according to their fraction cleaved value. Density was determined as 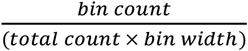 using base R for the calculations, as well as ggplot2(v3.5.1) and patchwork (v1.3.0) for the graphic. Python scripts used for the fraction cleaved analysis are available on GitLab.

Visualizing as a histogram revealed a bimodal distribution, with the two modes centered around the extreme levels of activity, either very high or very low FC. Most of the sequences (750) in the dataset were highly efficient self-cleaving ribozymes with FC values greater than 0.90 (**Table 1**). Of these,240 sequences were observed exclusively in the cleaved state (i.e. 100% cleaved). There were also another 40 sequences with a FC of 0.80-0.90, which also indicates efficient self-cleavage (**Table 1**). We verified high activity for four of these sequences by transcribing calculations, as well as ggplot2(v3.5.1) and patchwork (v1.3.0) for the graphic. Python scripts used for the fraction cleaved analysis are available on GitLab. them individually and quantifying their FC value by polyacrylamide gel electrophoresis (**Supplemental_Table_S2**). On the other extreme, there were 26 sequences that were either very inefficient or inactive ribozymes with a FC < 0.1, and seven more sequences with 0.1 < FC < 0.2 (**Table 1**). Five of these sequences were never observed in the self-cleaved state, but these sequences were also observed relatively few times (< 15 reads each) making it difficult to distinguish low efficiency ribozymes from completely inactive sequences. The distribution of FC observed is heavily skewed toward high-activity ribozymes. This further confirms that the bioinformatics approaches originally used to search for hairpin ribozymes were very efficient at discovering true, catalytically active ribozymes, and not just structurally similar sequences. The high activity of the ribozymes also supports the hypothesis that these ribozymes have a biological role in the circular RNA molecules where they have been predominantly found.

**Table 1.** Fraction cleaved values of hairpin ribozymes observed during deep-sequencing.

| Fraction Cleaved (FC) | Number of sequences |
| --- | --- |
| 0.0 – 0.1 | 26 |
| 0.1 – 0.2 | 7 |
| 0.2 – 0.3 | 3 |
| 0.3 – 0.4 | 1 |
| 0.4 – 0.5 | 9 |
| 0.5 – 0.6 | 2 |
| 0.6 – 0.7 | 9 |
| 0.7 – 0.8 | 8 |
| 0.8 – 0.9 | 40 |
| 0.9 – 1.0 | 750 |

[a] Counts are representative of the total number of sequences observed in either the cleaved or uncleaved state during analysis. See Experimental Details in the Supplementary Information for details. Python scripts used to determine fraction cleaved values are available on GitLab.

The RNA sequences with low activity are intriguing because it is challenging to understand why they would be found in natural samples if they are self-cleaving ribozymes that are very inefficient. We therefore set out to better understand the cause of decreased self-cleaving efficiency in the group of low activity ribozymes. A principal component analysis (PCA) was used to visualize all sequences and cluster them based on sequence similarity. This analysis only included regions corresponding to the consensus sequence positions (loops A and B and all four helices) so that length was constant and would not be the dominating source of sequence variation (**Figure 1A**). The data was colored according to their FC value to help identify any sequence clusters with low activity, which would indicate both similar sequence elements and similar lower activity. However, the analysis showed that sequences with low FC did not cluster and were distributed randomly throughout the first two dimensions of the principal component analysis (**Supplemental_Fig_S1)**. Since our library of sequences are diverse in length, we also checked if there was a correlation between length and FC, but did not find any significant correlation. This suggests that length alone is also not a simple indicator of low activity (**Supplemental_Fig_S2)**. These analyses confirmed that the low activity sequences do not have any obvious similarities.

The novel hairpin ribozymes were discovered through structural homology searches, which ensures that they can form the base pairs of the consensus hairpin secondary structure. However, it is possible that the less efficient ribozymes are less likely to adopt the consensus structure or form very unstable structures. To test this idea, we used Infernal to construct a hairpin ribozyme covariance model using the “extra-spruce-initial alignment” (Nawrocki & Eddy, 2013; Weinberg et al., 2021). We calibrated the model and matched our tested library of sequences to the covariance model to assign an E-value (Infernal *cmsearch* function). Higher E-values suggest a lower probability of a structural match to the consensus structure. All sequences had very low E-values (maximum of 5.9 × 10^−6^), but ranged over several orders of magnitude from each other. A correlation analysis did not indicate a significant correlation between FC and E-values (**Supplemental_Fig_S3**). Surprisingly, these analyses indicated that ribozymes with low activity do not consistently deviate from the consensus structure.

We noticed that the bimodal distribution of hairpin ribozyme Fraction Cleaved in **Figure 2** is very similar to the distribution of fitness effects (DFE) of random mutations that has been observed in RNA (Bendixsen et al., 2017). A bimodal DFE has also been observed from random mutations in proteins and genomes (Eyre-Walker & Keightley, 2007). For ribozymes, “fitness effects” of mutations are typically defined as the effect on activity relative to a wild-type ribozyme. The effect of individual random mutations in several RNA molecules has shown a bimodal distribution similar to our FC data, with mostly neutral or slightly deleterious mutations, and then a group of mutations with increasingly deleterious effects that are near or below the limit of detection. This similarity suggests that the RNA sequences in the metatranscriptomic data could be a random sampling of point mutations from catalytically efficient wild-type ribozymes. Because many of these ribozymes were found in viroid-like circular RNA molecules, it would be reasonable to expect elevated mutational rates due to error-prone RNA-dependent RNA Polymerase replication (Domingo & Perales, 2019; Lu et al., 2022). If the circular RNAs exist as a naturally occurring quasispecies containing numerous new, random mutations, the hairpin ribozyme sequences reported in the metatranscriptomic data would be a random sampling of these sequences. In addition, the metatranscriptomic sequencing itself could introduce mutations to the naturally occurring RNA through enzymatic mutations during library preparation, or sequencing errors. We note that these two sources of mutations are not mutually exclusive. Regardless of the source, random mutations could explain the distribution of FC observed in our data. Further investigations would be needed to test this idea, such as deeper sequencing of metatranscriptomic samples.

Several sequences in our library had rare sequence variations that nevertheless retained efficient self-cleavage in our experiments. For example, an adenosine nucleotide at position 10 (A10) is 90% conserved (**Figure 1A**), and it was previously reported that A10G mutation resulted in an 18-fold reduction in catalytic activity relative to the reference ribozyme without this mutation (Hampel & Tritz, 1989; Shippy et al., 1998). In our data, sequences with G10 were efficient self-cleaving ribozymes, with a FC = 0.975 ± 0.004 SE when averaged together (n = 55) (**Figure 3**). In addition, positions U41 and U42 in Loop B have also been previously shown to be important in facilitating cleavage and are 97% conserved (Berzal-Herranz et al., 1992) (**Figure 1A**). Two sequences in our data had a C41 nucleotide and one sequence had a G42 nucleotide. Both sequences with C41 were efficient self-cleaving ribozymes, with FC = 1.00 ± 0.000 SE and FC = 0.942 ± 0.002 SE (**Figure 3**). The sequence with G42 was still active, but less efficient with FC = 0.467 ± 0.129 SE (**Figure 3**). Notably, this sequence also contained a second “mutation” from the consensus structure at a 97% conserved residue (A43G), making it difficult to attribute the lower activity to either individual nucleotide change. At position G11 there is an uncommon G11A variation, which was expected to decrease self-cleavage because it changes a GC base pair to an AC (Weinberg et al., 2021). In our data, sequences with A11 were typically efficient self-cleaving ribozymes with an average FC = 0.917 ± 0.035 SE (n = 36), and only two sequences with FC ≤ 0.1 (**Figure 3**). Of the A11 sequences, 21 had a compensatory mutation at position -2, from a C to a U, and these sequences had high fraction cleaved as expected (FC = 0.864 ± 0.003 SE, n = 21). However, 15 sequences did not have this compensatory mutation, and still showed high fraction cleaved even with the AC mismatch (FC = 0.990 ± 0.058 SE, n = 15). Four additional ribozymes have guanine nucleobases at both positions 20 (R20) and 44 (N44), two positions that form a Watson-Crick base-pair in the crystal structure of the hairpin ribozyme from the minus strand of the tobacco ringspot virus satellite RNA ribozyme – a sequence not in our dataset (Rupert & Ferré-D’Amaré, 2001; Weinberg et al., 2021). However, the new consensus structure from the novel hairpin ribozymes does not indicate a conserved base-pair between position 20 and position 44 (**Figure 1A**). In our experiments, the four ribozymes with both G20 and G44 were active ribozymes with moderate to high FC values (**Figure 3, Supplemental_Table_S3**), confirming that a Watson-Crick base-pair is not required. It is possible that non-canonical base-pairing might be occurring between these positions. Combined, these results suggest that mutations at highly conserved residues may not be a good predictor of low ribozyme activity, and further structural investigations of new sequences are warranted. These results highlight a need for more accurate predictions of ribozyme activities, not just secondary structures, and for a better understanding of RNA sequence to function relationships in general.

**Figure 3.**
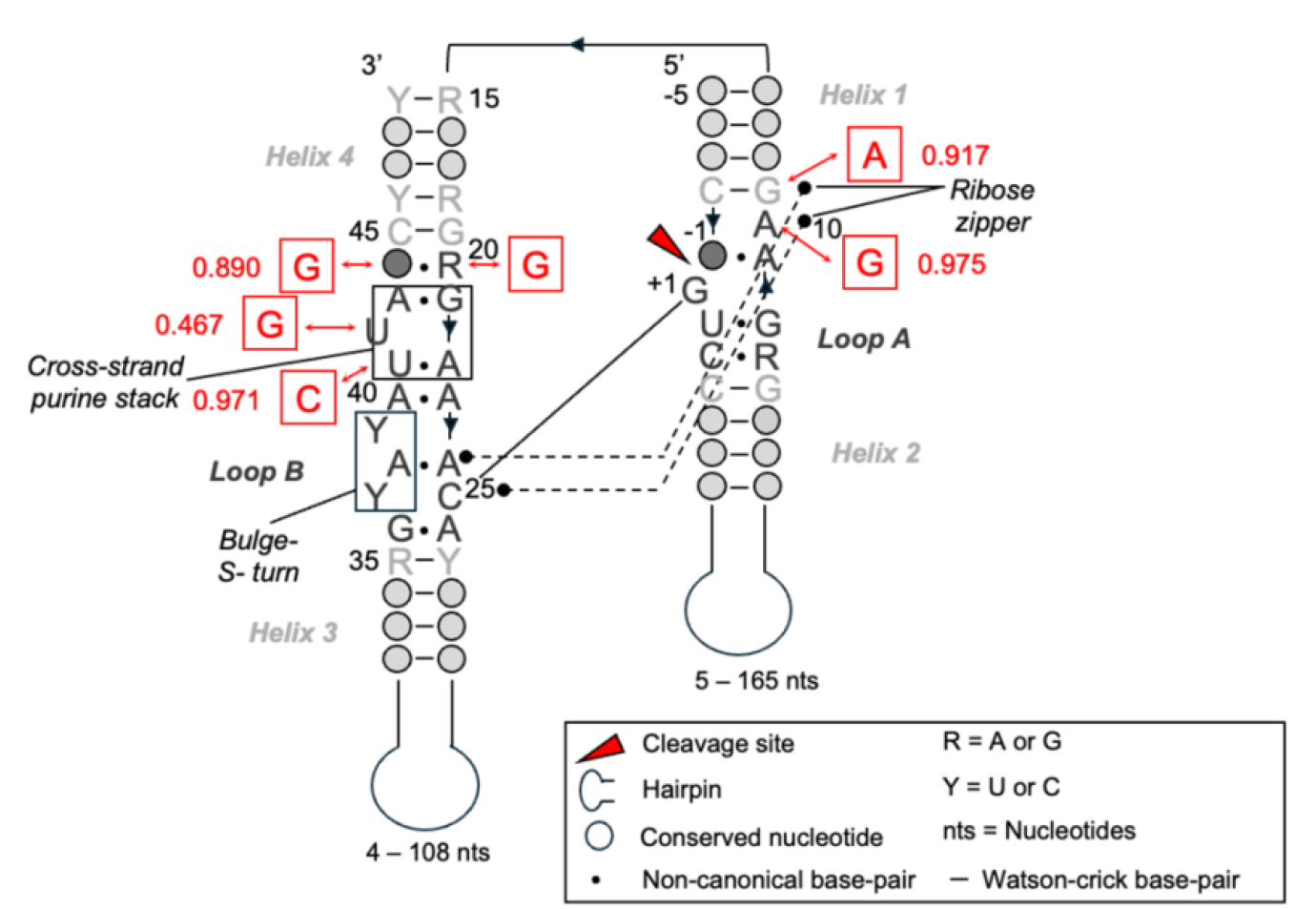
Sequence and structural characteristics of the consensus hairpin ribozyme based on the minus strand of the tobacco ringspot virus satellite RNA ribozyme crystal structure (Rupert & Ferré-D’Amaré, 2001). Relevant nucleotides are highlighted and boxed in red. Averaged fraction cleaved values for variant nucleotides are shown in red next to relevant positions. Python scripts used to determine nucleotide variants and fraction cleaved values are available on GitLab.

For many decades hairpin ribozymes were seen as rare in nature, which limited investigations into their biological roles. The discovery of hundreds of new examples of hairpin ribozymes motivates further investigation of this ribozyme class (Weinberg et al., 2021). What was known previously about the potential sequence diversity of this ribozyme was limited to deletion and mutagenesis studies (Berzal-Herranz et al., 1992; Roberts et al., 2023; Shippy et al., 1998), where the effects of mutations were only understood in the background of the four previously known genotypes. The data presented here confirmed that most of the newly discovered sequences are in fact very efficient self-cleaving ribozymes, including sequences with very different lengths, and rare sequence variation at highly conserved nucleotide positions. This data therefore expands our understanding of this ribozyme class, provides a new opportunity to evaluate sequence conservation, and also provides new sequences for engineering efforts (Hieronymus & Müller, 2021; Vauléon & Müller, 2003; Zhu et al., 2023). The four previously known variants of the hairpin ribozyme were all found in satellite RNAs of three viruses that infect species of agricultural significance (Buzayan et al., 1986; Kalvari et al., 2018; Kaper et al., 1988; Rubino et al., 1990). Their abundance in a metatranscriptomic dataset associated with “spruce litter” suggests that there might be important roles for other plant viruses in these natural environments, and further investigations are needed to uncover the biological roles and environmental consequences of these hairpin ribozymes.

## Methods

### PCR Amplification

All ssDNA oligonucleotides and the ssDNA library used in this study was made double-stranded for T7 transcription (**Supplemental_Table_S4**). PCR reactions included 10 µL 10 µM forward and reverse primers, 10 µL 10 µM DNA template, 10 µL 10X Standard Taq Reaction Buffer (New England Biolabs), 2 µL10 mM each dNTP mix (New England Biolabs), 2 µL Taq Polymerase (500U, Thermo Fischer) and 56 µL RNase-free water. PCR was performed using a Bio-Rad C1000 Touch Thermal Cycler. The thermal cycle was programed for 75 s at 95ºC for initial denaturation, followed by 34 cycles of 30 s at 95ºC for denaturation, 30 s at 53ºC for annealing, 30 s at 72ºC for extension, and 5 min at 72ºC for the final extension. PCR products were examined by 2% (w/v) sodium borate agarose gel electrophoresis at 200 V for 20 minutes in 1X sodium borate (10 mM sodium hydroxide, 36 mM boric acid, pH 8.5) running buffer. PCR products were premixed with 6X loading dye (NEB) that contained 1:200 GelRed® fluorescent DNA stain (Biotium). Gels were imaged on a SYNGENE G:Box under ethidium bromide settings. PCR products were cleaned and concentrated using the DNA Clean and Concentrator-5 kit from Zymo Research according to manufacturer instructions.

### Co-transcriptional self-cleavage assay reactions

The co-transcriptional self-cleavage reactions were carried out in triplicate by combining 10 µL 10X T7 transcription buffer (500 µL Tris pH 7.5, 50 µL 1M DTT, 20 µL 1M spermidine, 300 µL 1M MgCl_2_, 1 µL TRITON X-100 (0.1% v/v), 129 µL RNase-Free water), 4 µL T7 polymerase (In house high concentration > 20 U/µL), 2 µL 25 mM rNTP mix (New England Biolabs), 14 µL double-stranded DNA template (pool concentration 229.57 ng/µL), and 70 µL RNase-free water at 37ºC for 1 hour. The transcription and co-transcriptional self-cleavage reactions were quenched by adding 3.5 µL 0.5 M EDTA and placed on ice. In addition, 5 µL of TURBO DNase (1000U, Thermo Fischer) was added to the reactions and incubated for an additional 20 minutes at 37ºC to remove double-stranded DNA templates from the reaction.

### Reverse-transcription and Illumina Indexing reactions

Reverse transcription was carried out using a 5’ RACE protocol using phased template switching oligos. Reactions included 4 µL of 10 µM reverse transcription primer and 6 µL co-transcriptional self-cleavage product that was heated to 72ºC for 3 min and then cooled on ice. Reverse transcription was started by using 4 µL SMARTScribe 5X First Strand Buffer (TaKaRa), 2 µL 10 µM phased template switching oligo mix, 2 µL 20 mM DTT, 2 µL 20 mM DNTP MIX, and 2 µL SMARTScribe Reverse Transcriptase (200 units, TaKaRa). The mixture was incubated at 42ºC for 90 min and the reaction was stopped by heating at 72ºC for 15 min. Illumina adapter sequences and indexes were added in a reaction containing 3 µL of cDNA product, 12.5 µL 2X KAPA HiFi HotStart ReadyMix (KAPA Biosystems), 5 µL NGS Unique Dual Index Primers (IDT), and 4.5 µL RNase-Free water. Illumina indexing PCR was performed using a Bio-Rad C1000 Touch Thermal Cycler. The thermal cycle was programed for 75 s at 98ºC for initial denaturation, followed by 20 cycles of 10 s at 98ºC for denaturation, 30 s at 63ºC for annealing, 30 s at 72ºC for extension, and 1 min at 72ºC for the final extension. PCR products were examined by 2% (w/v) sodium borate agarose gel electrophoresis at 200 V for 20 minutes in 1X sodium borate (10 mM sodium hydroxide, 36 mM boric acid, pH 8.5) running buffer. PCR products were premixed with 6X loading dye (NEB) that contained 1:200 GelRed® fluorescent DNA stain (Biotium). Gels were imaged on a SYNGENE G:Box under ethidium bromide settings. PCR products were cleaned and concentrated using the DNA Clean and Concentrator-5 kit from Zymo Research according to the manufacturer instructions.

### High-throughput sequencing

Indexed PCR products for all replicates were pooled together at equimolar concentrations based on a DeNovix OFX Fluorometer standard calibration curve using the Biotium AccuClear® Ultra High Sensitivity dsDNA Quantification Kit. Paired-end sequencing reads were obtained for the pooled libraries using an Illumina MiSeq v2 micro, 300 cycle run type (Genomics and Cell Characterization Core Facility, University of Oregon).

### Data analysis and visualization

Paired-end sequencing reads were joined using FLASh (v1.2.11), allowing ‘outies’ due to overlapping reads. The joined sequencing reads were then analyzed using a custom python jupyter lab notebook. Python (v3.9.6) was used to filter, extract and analyze read counts. R (v4.4.1) was used for data visualization and Pearson R coefficient analysis. Infernal (v1.1.5) was used for consensus structure analysis using a developed covariation model. Prior to all data analysis, sequences were trimmed by removing low quality bases that had a Phred quality score less than or equal to 20 were removed with a custom python script using the BioPython SeqIO (v1.43) module. Reads were then extracted based on regular expression patterns and matched to their respective reference sequence within the tested library. The cleavage status and number of total occurrences was recorded for each genotype so that the fraction cleaved could be quantified. The fraction cleaved value (mean probability, n = 3) was determined as described in the main text using the total number of cleaved and uncleaved read counts for each individual ribozyme. Standard error was calculated as:

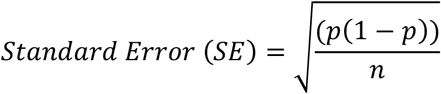

Where p is the mean probability fraction cleaved value of a ribozyme and n represents the total number of observations for that ribozyme sequence. 95% confidence intervals were determined by multiplying the calculated standard error value by 1.96, which represents z-score for 95% of the area under a normal distribution curve. The Infernal covariation model was built with the following command lines: *cmbuild* was used to construct the covariance model using the “extra-spruce-initial alignment” from the previous bioinformatics approach*. Next *cmcalibrate* was employed to calibrate the previously constructed model so E-values could be determined.

Lastly, *cmsearch* was used to predict E-values based on an ungapped fasta file of the hairpin ribozymes used during our high-throughput based assay. We note that 17 sequences expected to be in the data were never observed in either the cleaved or uncleaved state, which could be due to “drop-outs” in the DNA synthesis, and/or low efficiency in any of the assay steps (transcription, reverse-transcription, PCR). Furthermore, 13 pairs of sequences were found to have identical sequences except at their cleaved 5-nt 5’ end. As a result, this prevented their evaluation by deep-sequencing and was subsequently removed from analysis. Code used in our analysis was made with the assistance of ChatGPT(v2).

### Individual ribozyme assays

Individual ribozyme sequences were ordered as ssDNA template oligos (IDT) and were made double-stranded by PCR, and underwent co-transcriptional self-cleavage reactions as previously described (**Supplemental_Table_S4**). To better visualize uncleaved and cleaved fragments, DNA template concentrations for co-transcriptional self-cleavage reactions were increased to 3.34 ng/µL. The transcribed RNAs were separated by 10% denaturing PAGE (8M urea) and stained using 1:200 GelRed (prepared using 10,000X GelRed®, Biotium). The gels were imaged using a SYNGENE G:Box under ethidium bromide settings. Fraction cleaved values were determined by gel densitometry using GelAnalyzer (v23.1.1) software under default settings.

## Data Availability

Custom python scripts and FastQ files used in the processing and analysis of the data can be found on GitLab (https://gitlab.com/bsu/HaydenLab_KM/hairpinribozyme).

## Acknowledgments

We would like to acknowledge Dr. Allison Simler-Williamson for her help and support with the statistical analysis of the hairpin ribozyme dataset.

